# Blood transcriptome analysis suggests an indirect molecular association of early life adversities and adult social anxiety disorder by immune-related signal transduction

**DOI:** 10.1101/2022.12.22.521187

**Authors:** Susanne Edelmann, Ariane Wiegand, Thomas Hentrich, Sarah Pasche, Julia Schulze-Hentrich, Matthias H. J. Munk, Andreas J. Fallgatter, Benjamin Kreifelts, Vanessa Nieratschker

**Author notes:** **Correspondence:** Vanessa Nieratschker.

## Abstract

Social anxiety disorder (SAD) is a psychiatric disorder characterized by severe fear in social situations and avoidance of these. Multiple genetic as well as environmental factors contribute to the etiopathology of SAD. One of the main risk factors for SAD is stress, especially during early periods of life (early life adversity; ELA). ELA leads to structural and regulatory alterations contributing to disease vulnerability. This includes the dysregulation of the immune response. However, the molecular link between ELA and the risk for SAD in adulthood remain largely unclear. Evidence is emerging that long-lasting changes of gene expression patterns play an important role in the biological mechanisms linking ELA and SAD.

Therefore, we performed a transcriptome study of SAD and ELA using RNA sequencing. Analyzing differential gene expression, 13 significantly differentially expressed genes (DEGs) were identified with respect to SAD whilst no significant differences in expression were identified with respect to ELA. The most significantly expressed gene was *MAPK3* being upregulated in the SAD group compared to control individuals. In contrary, weighted gene co-expression network analyses (*WGCNA*) identified only modules significantly associated with ELA, not with SAD. Furthermore, analyzing interaction networks of the genes from the ELA-associated modules and the SAD-related MAPK3) revealed complex interactions of those genes. Gene functional enrichment analyses indicate a role of signal transduction pathways as well as inflammatory responses supporting an involvement of the immune system in the association of ELA and SAD.

In conclusion, we did not identify a direct molecular link between ELA and adult SAD by transcriptional changes. However, our data indicate an indirect association of ELA and SAD mediated by the interaction of genes involved in immune-related signal transduction.

## 1. Introduction

Anxiety disorders are common and highly comorbid with other psychiatric disorders [1]. A distinctive form is Social Anxiety Disorder (SAD) with an estimated worldwide lifetime prevalence of 4 % [2]. SAD is described by severe fear and avoidance behavior in social situations such as fear of being the center of attention or fear of negative social evaluation which can have a detrimental impact on daily life [3]. The etiology of SAD is influenced by genetic [4, 5] as well as environmental factors. One of the most relevant environmental influences is stress. Stressful experiences in critical periods of life, especially during childhood and adolescence, can lead to structural and regulatory alterations—such as disturbed programming of the hypothalamic-pituitary-adrenal (HPA) axis—contributing to disease vulnerability [6-8]. Furthermore, dysregulation of the—inflammatory—immune response through childhood stress exposure can affect brain development, cognition, stress reactivity and resilience and, hence, the risk for psychopathology later in life [9-12]. Early life adversity (ELA) therefore represents one of the main environmental factors contributing to an increased risk for SAD [13, 14]. However, the molecular link between an early stressor, such as adverse events during childhood, and the risk for SAD in adulthood remains unclear.

Changes of gene expression patterns following ELA have been identified in different organisms [15-17]. In humans, monocytes of individuals exposed to early childhood maltreatment showed altered HPA axis responses to stress, evidenced by lower blood adreno-corticotropic hormone and cortisol levels. Moreover, the analysis of transcriptome-wide gene expression patterns in the same samples showed that stress-responsive transcripts were enriched for genes involved in cytokine- and inflammation-related pathways [18]. In addition, co-expression network analysis identified an association of ELA with inflammation-related pathways [19]. Furthermore, RNA sequencing (RNA-seq) in brain tissue revealed enrichment of differentially expressed genes in immune and GTPase function in individuals with a history of ELA as compared to control individuals without the experience of ELA [20].

Aberrant gene expression patterns of various genes, with some of them involved in the immune system, have also been identified in humans across different social environments such as social isolation or low socioeconomic status [21-23]. Moreover, the expression of genes involved in immune response as well as transcriptional regulation and cell proliferation has been shown to be sensitive to social regulation (more precisely, the level of loneliness, [24]). Therefore, not only ELA, but also acute social stress is likely to impact gene expression in humans as it has already been proven in mice, in which the vascular system and inflammatory pathways were mainly affected [25]. Furthermore, several studies have indicated an association between expression changes of diverse genes in mouse brain and social fear [26] as well as anxiety [27, 28]. In humans, an investigation of the blood transcriptome has suggested altered immune functions in generalized anxiety disorder [29]. In addition, expression differences of α-endomannosidase (MANEA) are associated with SAD and panic disorder in human blood [30]. Moreover, RNA-seq [31] has identified higher *ITM2B* gene expression levels associated with higher anxiety scores in a cohort of 25 monozygotic (MZ) twins, which has been validated in a second cohort of 22 MZ twins [31].

A molecular link between ELA and the sensitivity to social stress on the transcriptome level has been shown in mouse brain tissue, where distinct transcriptional patterns depending on ELA in socially stressed adult mice have been revealed: Several genes have been identified as differentially expressed in mice with ELA and social stress in adulthood compared to controls without ELA. Interestingly, their gene expression levels have not been altered when exposed to either ELA or adult social stress alone [17]. However, in humans the association between transcriptional changes induced by ELA and adult SAD still remain elusive.

In the current study we aimed to identify gene expression patterns associated with SAD, ELA and their interaction on a transcriptome-wide level. We performed RNA-seq in whole blood of individuals with SAD with high or low levels of ELA and control individuals with high or low levels of ELA, respectively, to explore gene expression differences and transcriptional networks. We identified genes differentially expressed in the context of SAD and clusters of co-expressed genes associated with emotional ELA. Those genes were shown to closely interact. Our results indicate an indirect molecular link between emotional ELA and SAD mediated by the interaction of genes involved in immune-related signal transduction.

## 2. Material and methods

### Study population

In total, 160 participants of Caucasian descent between 19-50 years of age took part in the study. All participants were assessed using the Structured Clinical Interview for DSM-IV (SCID) and 70 participants were found to be suffering from SAD as a primary diagnosis. The severity of social anxiety was evaluated using the Liebowitz Social Anxiety Scale (LSAS, [32]). ELA was assessed using the Childhood Trauma Questionnaire (CTQ) that measures five dimensions (further referred to as subscales) of maltreatment: emotional and physical neglect and emotional, physical and sexual abuse [33, 34]. Participants with at least a moderate score in one of the five categories were classified as participants with high levels of ELA [34, 35]. Thus, four groups emerged: 1) control participants without SAD and low levels of ELA (n=62), 2) control participants without SAD and high levels of ELA (n=27), 3) participants suffering from SAD with low levels of ELA (n=43), and 4) participants suffering from SAD with high levels of ELA (n=27). All participants gave written informed consent to the experimental procedure prior to inclusion in the study. The study was performed in accordance with the Declaration of Helsinki and approved by the University of Tübingen local ethics committee.

### RNA extraction, library preparation and sequencing

Total RNA from whole blood stored in PAXgene Blood RNA tubes was extracted using the PaxGene Blood miRNA kit (Qiagen, Hilden, Germany). Quality of RNA was assessed using a Bioanalyzer (Agilent, Santa Clara, USA). Only samples with an RNA integrity number (RIN) of 7 and higher were used for sequencing library preparation. Libraries for 3**′** RNA-seq were prepared using the 3**′** method by Lexogen [36] as used in the NGS Competence Center Tübingen (NCCT) where both library preparation and sequencing in randomized batches was performed. First strand synthesis of polyA-tailed RNA from total RNA using oligo dT primers was followed by degradation of the RNA template, second strand synthesis with random primers containing 5**′** Illumina-compatible linker sequences and amplification using random primers that add barcodes and cluster generation sequences [36]. The libraries were sequenced on the NCCT Nova sequencing platform at a depth of about 10 million reads with 100 bp in length.

### RNA-seq reads preprocessing

Read preprocessing was performed using the Lexogen pipeline [37] implementing the *bbduk* tool from the BBTools suite (https://sourceforge.net/projects/bbmap/) for quality trimming and the STAR aligner [38] for mapping to the reference genome (vGRCh38.104). Principle component analysis (PCA) was used to detect sample outliers by *DESeq2* [39]. The *R* package *OUTRIDER* [40] was used to identify gene count outliers that were excluded from further analyses (Table S1).

To control for the effect of blood cell type composition variability on gene expression, blood cell type proportions were estimated using the *granulator* package in *R* using TPM (transcripts per million) normalized counts. Benchmarking in *granulator* was performed using reference cell type counts of a subset of the cohort (Fig. S1). The R package *variancePartition* [41] was used to calculate the variance explained by differential cell type composition and covariates. The package implements a linear mixed model method to characterize the contribution of selected variables to transcriptional variability. As deconvolution results showed a minor contribution of most cell types to the variance between the samples (Fig. S2), we used an adjustment approach of the gene counts to all cell type ratios resulting from the deconvolution approach based on a linear model adapted from Jones *et al*. [42] instead of using the cell type ratios as covariates in the later analyses.

### Data analyses

#### Demographic and Clinical Information

Normality of data was tested using Shapiro-Wilk test. The test revealed non-normal distributions for all variables (Table S2). Therefore, the comparison of the trait medians between the independent groups was performed using the Wilcoxon Mann-Whitney rank sum test.

#### Differential Gene Expression (DGE)

Differential gene expression analysis was performed using *DESeq2* [39], which analyzes differences in gene expression based on a negative binomial generalized linear model. Genes with low counts were removed and only those with at least 20 counts in all samples were kept, as huge on/off changes were not expected due to the research question. A linear model with the factors of interest SAD, ELA and covariates age and sex was fitted. The false discovery rate (FDR) to adjust the *p* value to multiple correction) was set at 0.1. Results were filtered for differentially expressed (DE) genes with an absolute log2 fold-change larger than 0.3.

#### Weighted Gene Co-expression Network Analysis (WGCNA)

Scale-free co-expression networks were constructed using the *R* package *WGCNA* that defines modules using a dynamic tree-cutting algorithm based on hierarchical clustering of expression values (minimum module size = 100, cutting height = 0.99). *WGCNA* was performed using filtered (≥ 20 counts per sample) and variance stabilized count data (generated from the read count matrix using *DESeq2*’s *getVarianceStabilizedData* function). The network was constructed at a soft power of 10 at which the scale-free topology fit index reached 0.9. The module eigenvalue was used to perform the correlation analysis with the variables (i.e. questionnaire scores of LSAS and its subcategories as well as CTQ and its subcategories; covariates age and sex) with each whole module. Modules additionally significantly correlating with sex and age were discarded from further analyses.

#### Gene Functional Enrichment Analysis

Gene list functional enrichment analysis was performed using the *R* package *gProfiler2* [43, 44] by using the *Gene Ontology* (GO) resource (vOBO 1.4, [45, 46]), the Kyoto Encyclopedia Genes and Genomes (*KEGG*) pathways database (v103.0, [47]) and *Reactome* database (v81, [48]). Terms with FDR-corrected *p* values of < 0.05 were considered significantly enriched within modules.

#### Network Analysis and Visualization

*MAPK3*, the top hit of the DGE analysis, was imported into the online Search Tool for the Retrieval of Interacting Genes/Proteins (STRING) database v11.5 (http://string-db.org; [49]) for known and predicted protein-protein interactions (ppi). We used the following conditions for network generation: medium confidence (0.4), maximum 50 interactors for the first shell and 10 for the second shell.

The interactome of MAPK3 together with all genes of significant *WGCNA* modules (in total 1815) was generated using the STRING database (v11.5) starting with a *full network* (edges indicating both functional and physical protein associations) and then filtering for interaction scores > 0.9, thereby increasing confidence. For the final interactome, all direct neighbors of MAPK3 were selected. The interactome was visualized using Cytoscape (v3.9.1., [50]).

## 3. Results

### Demographic and Clinical Information

Table 1 shows the sample characteristics with respect to the four groups emerging from the factors SAD and ELA in more detail. While there was neither a significant group difference in age (Wilcoxon, n = 159, W = 3370, *p* = 0.38) nor sex (Pearson’s Chi-square test, χ^2^ = 1.62, *p* = 0.20) with respect to SAD, significant differences in age (Wilcoxon test, n = 159, W = 3398, *p* = 0.040), but not sex (Pearson’s Chi-square test, χ^2^ = 0.05, *p* = 0.82) emerged with respect to ELA. Additionally, Levene’s test revealed variance heterogeneity of the age data among the ELA groups (DF = 1, F = 14.814, *p* = 0.000).

**Table 1:** Sample characteristics for the four groups emerging from the factors SAD and ELA. Mean ± standard deviation. SAD = Social Anxiety Disorder, ELA = Early Life Adversity, LSAS = Liebowitz Social Anxiety Scale, CTQ = Childhood Trauma Questionnaire.

|  | group |  |  |  |
| --- | --- | --- | --- | --- |
|  | SAD |  | no SAD |  |
|  | high ELA | low ELA | high ELA | low ELA |
| **n** | 27 | 43 | 27 | 62 |
| **LSAS total score** | 75.8 (± 28.3) | 68.9 (± 25.3) | 23.6 (± 17.2) | 14.8 (± 14.1) |
| **CTQ total score** | 56.6 (± 13.8) | 32.7 (± 5.2) | 48.5 (± 10.3) | 29.7 (± 4.6) |
| **Sex** | ♀18 ♂9 | ♀32 ♂11 | ♀17 ♂10 | ♀38 ♂24 |
| **Age [years]** | 29 (± 8) | 24 (± 6) | 27 (± 8) | 25 (± 3) |

The total score of the CTQ of our cohort was not significantly different with respect to sex (Wilcoxon test, n = 159, W = 2373.5, *p* = 0.093), but it correlated positively with age (r = 0.117, *p* < 0.001). Nevertheless, we included age (in addition to sex) as covariate in the DGE analysis and tested each candidate gene expression count for correlation with age to exclude age effects on the expression of the respective gene (Fig. S3). For the total score of the LSAS, there was no significant difference with respect to sex (Wilcoxon test, n = 159, W = 3337, *p* = 0.068) and no correlation with age (r = −0.001, *p* = 0.83). Finally, there was a significant correlation between the total scores of the LSAS and CTQ (r = 0.104, *p* < 0.001). This correlation was mainly due to the highly significant correlation of the emotional CTQ subscales emotional abuse (r = 0.107, *p* < 0.001, n = 29) and neglect (r = 0.092, *p* < 0.001, n = 35) with the LSAS score (Fig. S4), whereas the other subscales of the CTQ did not or less significantly correlate with the LSAS total score (physical abuse: r = 0.018, *p* = 0.05; sexual abuse: r = 0.018, *p* = 0.06; physical neglect: r = 0.004, *p* = 0.010). Importantly, in our cohort we have eight cases of physical and five cases of sexual abuse only (Table S3), which needs to be kept in mind when interpreting the results.

### Differential Gene Expression with respect to SAD, but not ELA

Investigating gene expression of all participants, visual inspection of the PCA revealed no obvious grouping of samples (Fig S5). This is in line with rather subtle gene expression changes that one may expect in blood in the context of mental disorders [51, 52]. Analyzing differential gene expression using *DESeq2* [39], 13 significantly (FDR-corrected *p* ≤ 0.1) differentially expressed genes (DEGs) were identified which had a |l2fc| ≥ 0.3 with respect to SAD (Fig. 1A, Table S4), whilst no significant differences in expression were identified with respect to ELA (Table S5).

**Fig. 1:**
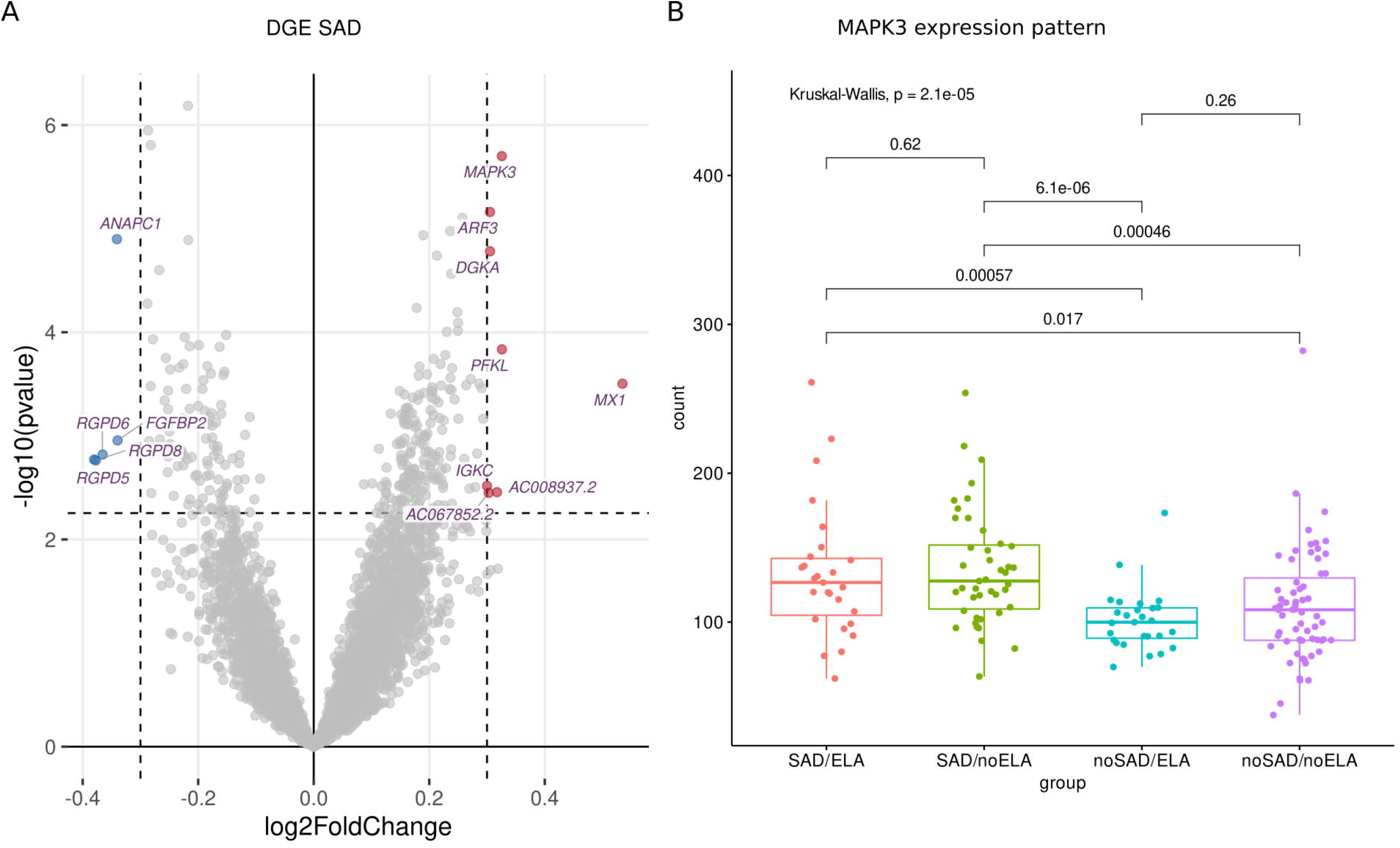
DGE analysis between SAD and control group. **A** Volcano plot displaying the significantly (FDR-corrected *p* ≥ 0.1) differentially (|l2fc| ≥ 0.3) expressed genes with down-regulated genes marked in blue and up-regulated genes marked in red. **B** The expression patterns of the most significantly differentially expressed gene *MAPK3* displayed in gene counts with respect to SAD and ELA show increased gene counts in the SAD groups without an influence of ELA. Wilcoxon rank sum test was applied and p values were adjusted for multiple testing using Benjamini-Hochberg correction. Kruskal-Wallis-Test additionally shows significantly different *MAPK3* expression between the SAD/ no SAD (with respect to ELA) groups.

Visualizing the count distribution of all SAD associated DEGs, eight of the candidates exhibited an expression pattern rather caused by extreme values (such as outliers) or other effects than being a true DEG (Fig. S6). Therefore, we excluded these genes from further analyses (marked in Table S4). The remaining DEGs include (in order of significance) *MAPK3* (Mitogen-Activated Protein Kinase 3), *ANAPC1* (Anaphase Promoting Complex Subunit 1), *PFKL* (Phosphofructokinase, Liver Type), *FGFBP2* (Fibroblast Growth Factor Binding Protein 2) and *AC008937*.*2* (long non-coding (lnc) RNA, Fig S7). The most significantly expressed gene *MAPK3* (FDR-corrected *p* = 0.003, l2fc = 0.33) was upregulated in the SAD group compared to control individuals (Fig. 1B). Fig. 1B shows the counts of *MAPK3* for each experimental group revealing that there is no ELA specific expression pattern within the SAD group. SAD groups with respect to ELA displayed an equally high *MAPK3* count (SAD/ELA: adj. mean gene count = 132.59 ± 44.71, SAD/no ELA: adj. mean gene count = 135.62 ± 38.59), whereas the groups of individuals without SAD showed a significantly lower *MAPK3* mean count, no matter whether ELA levels were high or low (no SAD/ELA: adj. mean gene count = 101.21 ± 20.60, no SAD/no ELA: adj. mean gene count = 110.49 ± 38.06).

In addition to the comparison of ELA groups, the CTQ subscales were classified according to Bernstein and Fink [53], with a score moderate and higher indicating the respective trauma (Table S3). DGE analysis was carried out for each CTQ subscale (Table S5). Furthermore, we analyzed DGE in the SAD group only with respect to ELA and each subscale as well as in the ELA group with respect to SAD to examine the potential transcriptomic association of ELA and SAD (Table S5). The subscale DGE analyses resulted in two genes differentially expressed in individuals with or without the experience of childhood sexual abuse in the entire cohort (Table S6) and 197 genes with respect to physical abuse within the SAD group only (Table S7). However, as there were only five cases of sexual abuse in the entire cohort and four cases of physical abuse in the SAD group only (Table S3), the DGE analysis in those subgroups has to be interpreted with caution. In summary, there were no relevant significant DEGs identified for ELA as well as the CTQ subscales neither in the entire cohort nor in the SAD group only.

### Gene Co-expression Clusters correlating with emotional ELA

*WGCNA* uses a validated principle called guilt by association, which relies on the assumption that associated or interacting genes share expression patterns and are likely to function together. Therefore, *WGCNA* was performed on gene counts matching the same filters as for the DGE analysis and a soft threshold power of 10 to identify ELA and/or SAD specific gene co-expression (for more details on the analysis, see Fig. S8). We identified 11 gene co-expression modules correlating with any of the variables available for the cohort (Table S8) with sizes ranging from 73 to 1750 genes. 1559 genes were assigned as not correlated (module grey).

Interestingly, whereas the DGE analysis revealed only associations of gene expression and SAD, the WGCNA resulted only in modules significantly correlated with ELA but not SAD or LSAS scores, respectively. In more detail, the modules red (Table S9), greenyellow (Table S10) and turquoise (Table S11) were significantly correlated with the ELA groups (turquoise), the subscales emotional abuse and emotional neglect as well as total CTQ score (red and greenyellow), respectively, but not with any of the non-disease-related variables (age, sex, size and weight, Table 2). None of the ELA co-expression cluster top hub genes (Table S12) overlapped with SAD DEGs. Furthermore, *MAPK3* was found in the grey module containing the genes not correlated with any variable. The other relevant DEGs were found in the following co-expression modules: the green module that was not significantly correlated with any variable (Fig. S9) included *ANAPC1* and *FGFBP2. PFKL* was found in the blue module which is associated with (amongst others) ELA, sex and size (Fig. S9). Finally, *AC008937*.*2* was co-expressed with genes in the red module.

**Table 2:**
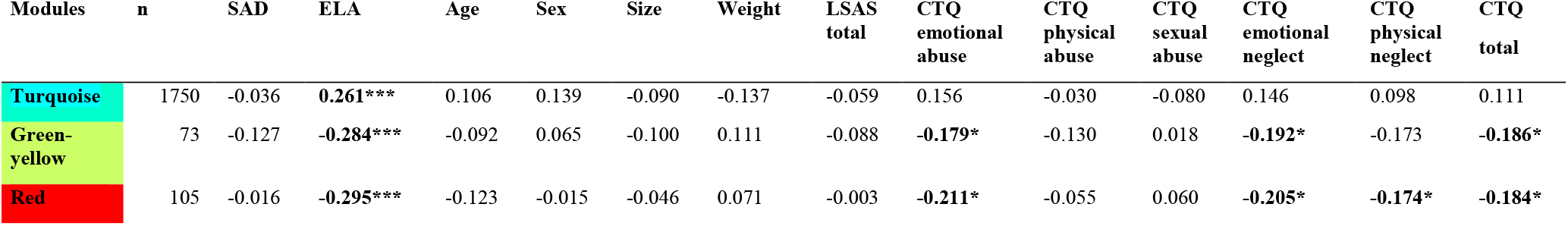
*WGCNA* module gene count and correlation coefficients for each variable. *** *p* ≤ 0.01, * *p* ≤ 0.05.

### Gene functional enrichment analysis reveals relevance of signal transduction pathways and immune system

A protein-protein-interaction (ppi) network of MAPK3 was generated using the STRING database to find pathways in which co-expressed genes within ELA-specific WGCNA modules and MAPK3-associated with SAD potentially interact. A functional enrichment analysis was performed to examine the enrichment of annotated terms within the three modules significantly correlating with ELA and/or the respective CTQ (sub-)scales modules (turquoise, red and greenyellow) and the MAPK3 ppi network. The MAPK3 ppi network was enriched mainly for MAPK signal transduction pathways and NTRK (neurotrophin receptor) signaling. The red and the turquoise *WGCNA* modules were enriched for cellular structural processes/compartments (Fig. 2A and B). The greenyellow co-expression cluster contained genes particularly involved in immune-related pathways (especially interleukin regulation and production) and JAK-STAT signaling (Fig. 2C).

**Fig. 2:**
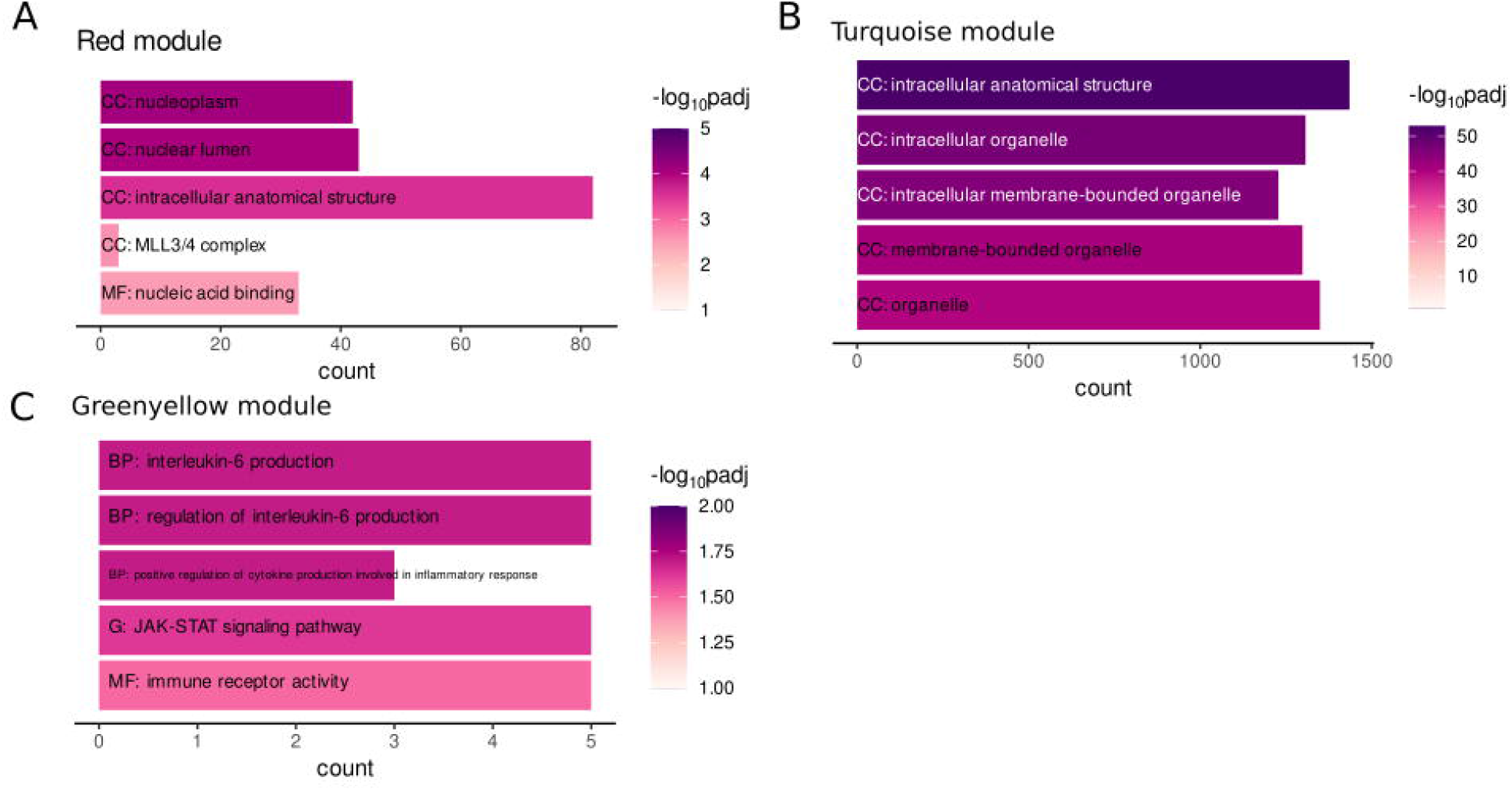
**Gene functional enrichment** of the **A** *WGCNA* module red, **B** *WGCNA* module greenyellow and **C** *WGCNA* module turquoise. Significance values are color-coded. Abbreviations indicate the database with G: KEGG database, BP: Biological process (GO term), CC: Cellular component (GO term) and MF: Molecular function (GO term).

The enrichment analysis did not reveal any shared or overlapping pathways between the modules, which does not indicate a direct molecular mediation of ELA on adult SAD by one single process.

### Network Analysis identifies common Genes between SAD-related MAPK3 and ELA associated Co-Expression Modules

The gene lists of each co-expression module correlating with ELA and/or CTQ (sub)scales, i.e. of the turquoise, red and greenyellow modules, were compared with the genes contained in the MAPK3 ppi network that plays a role in SAD to identify overlapping and thus potentially interacting genes. We identified *PTPN7* (Tyrosine-protein phosphatase non-receptor type 7) as a gene present in the MAPK3 ppi network and the red module (Table S13). PTPN7 is a member of the phosphatase family and a known negative regulator of MAPK signal cascade activation [54]. The turquoise module and the MAPK3 ppi network share a set of 20 genes (Table S13). Gene set enrichment analysis identified an involvement of most of those genes especially in signal transduction (MAPK, NTRK, neurotrophin, Fig. S10). The hub gene of the greenyellow module *STAT3* (Signal transducer and activator of transcription 3) as well as *RAF1* (RAF Proto-Oncogene Serine/Threonine-Protein Kinase) were also found in the MAPK3 ppi network (Table S13). RAF1 activation initiates a mitogen-activated protein kinase cascade and is in part regulated by cytokine signaling [55] and STAT3 mediates cellular responses to interleukins and other growth factors [56-62] as well as inflammatory responses by regulating differentiation of naive CD4+ T-cells into T-helper Th17 or regulatory T-cells [63]. Therefore, both genes are involved in the immune response and are linked to the mitogen-activated signaling cascade [55, 64, 65], where MAPK3 plays a central role[55]. Therefore, an interaction of STAT3, RAF1 and MAPK3 in immune signaling is likely.

To verify the interaction of MAPK3 and genes from the ELA-correlated modules, we used the STRING database to extract information on interaction scores of the 1815 genes (MAPK3 + genes from the three modules). After removing all genes with an interaction score ≤ 0.9 indicating the highest confidence, 51 genes remained. We generated an interactome of those genes (Fig. 3). The interactome highlighted the interaction of MAPK3 and the beforementioned STAT3, RAF1 and PTPN7. However, the interactome also revealed complex interrelations between the genes with several to many interaction partners of each gene (Fig. 3).

**Fig. 3:**
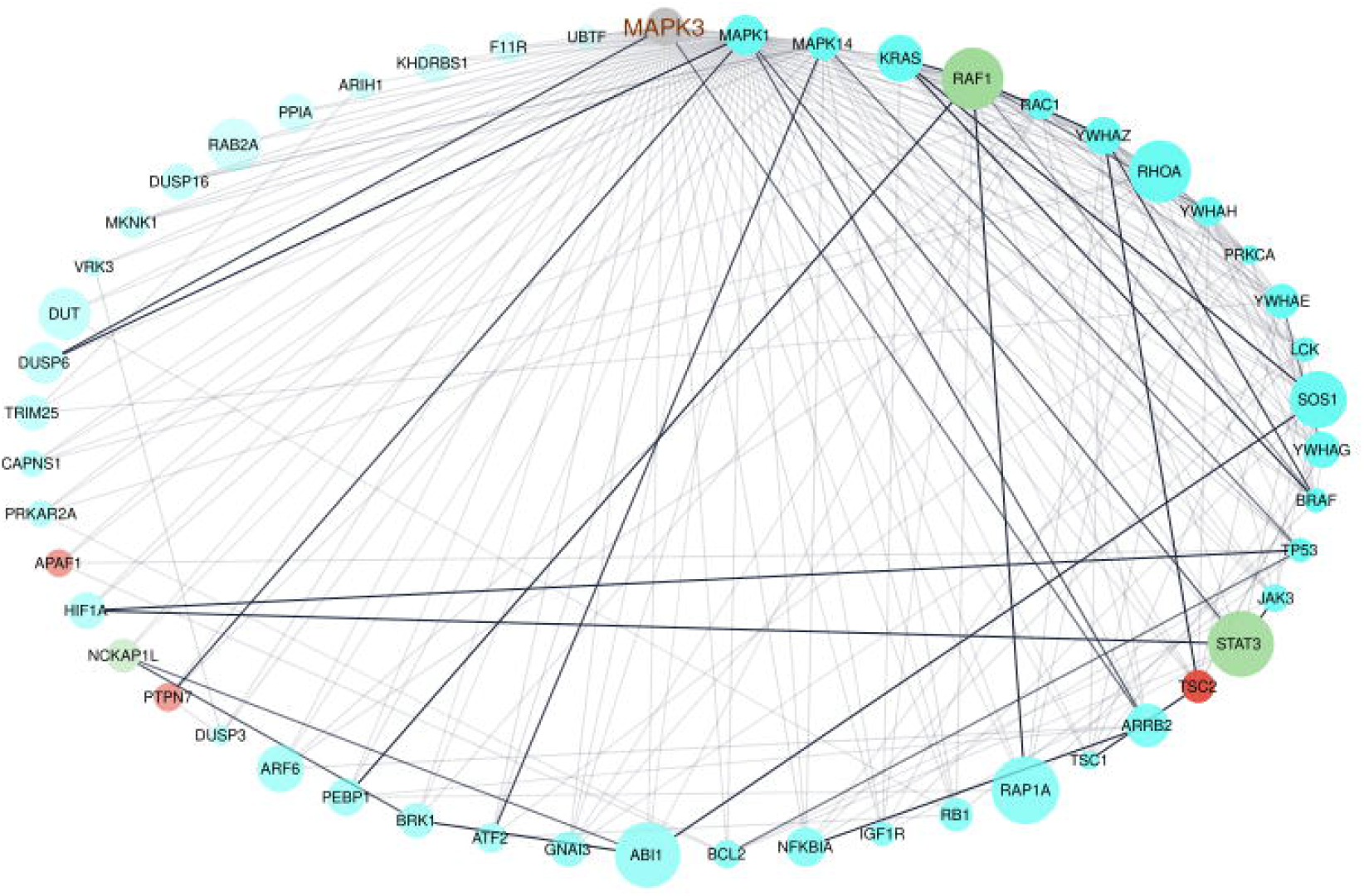
Network visualization of the interactome of MAPK3 and the genes from the turquoise, red and greenyellow module. The nodes were colored by module, node size displays module membership score from *WGCNA*, the node sort and node transparency were set by STRING degree and the edge transparency was set by STRING score.

The functional enrichment of the gene list was performed by using the *Reactome* database only. This enabled focusing on interaction of the genes to form a biologically relevant network as the *Reactome* groups entities participating in reactions. The analysis revealed mainly enrichment of the genes in terms related to immune-related signaling (Fig. 4, Signaling by Receptor Tyrosine Kinase, Signaling by NTRKs, etc.). However, the most significantly enriched pathway was Signal Transduction (Fig. 4).

**Fig. 4:**
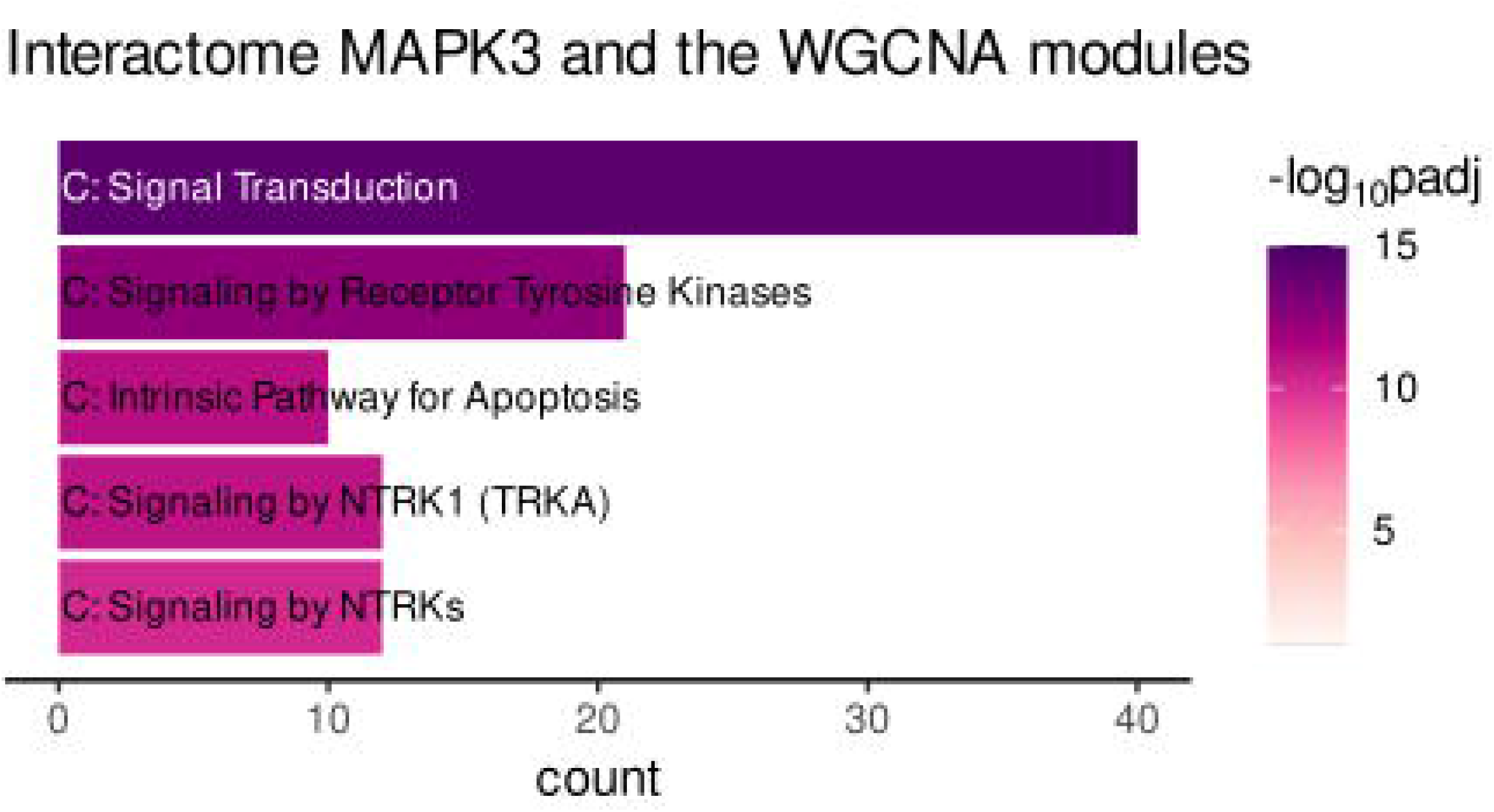
Gene functional enrichment of the *MAPK3* interactome associated with genes of the turquoise, red and greenyellow module. Significance values are color-coded.

## 4. Discussion

Transcriptome analyses have become highly relevant over recent years to investigate the molecular basis of psychiatric disorders. In particular, RNA-seq has been widely used to analyze psychiatric disorders and interrelations [66-68]. In the study presented here, we focused on social anxiety disorder and the molecular connection with a potential environmental trigger—early life adversity.

DGE analysis revealed genes associated with SAD, with *MAPK3* being the most significantly upregulated in individuals with SAD compared to control individuals. No DEGs were identified between individuals with and without a history of ELA. MAPK3 is a serine/threonine kinase which acts as an essential component of the MAP kinase signal transduction pathway. MAPK1/ERK2 and MAPK3/ERK1 play an important role in the MAPK/ERK cascade (extracellular signal-regulated kinase-dependent cascade). Depending on the cellular context, the MAPK/ERK cascade mediates diverse biological functions such as cell growth, adhesion, survival and differentiation through the regulation of transcription, translation and cytoskeletal rearrangements. The MAPK/ERK cascade also plays a role in initiation and regulation of meiosis, mitosis, and postmitotic functions in differentiated cells by phosphorylating a number of transcription factors that, for example, promote breast cancer [69]. Interestingly, another finding was the differential expression of the lncRNA AC0008937.2 in the context of SAD, which—as an antisense lncRNA to *MAP3K1*—has a potentially regulative role [70] in the MAPK signaling cascade. Another important gene that in turn is regulated by MAPK signaling is the brain-derived neurotrophic factor (BDNF) [71, 72]. The neurotrophin BDNF is essential for dendritic development in peripheral and central nervous system and regulates dendritic growth [73]. Moreover, BDNF level changes in serum are associated with a variety of anxiety disorders [74, 75]. Furthermore, BDNF protein levels were shown to be reduced in brain tissue of rats with a history of ELA which were subsequently exposed to stress [76]. However, in our data derived from whole blood we did not identify a SAD-specific expression pattern of *BDNF*. MAPK3 may function as a transporter in blood to regulate the expression of *BDNF* in brain tissue which in turn may lead to altered structural brain plasticity playing a role in SAD. Therefore, the analysis of *BDNF* expression levels would be of interest in different brain areas in the context of SAD. *MAPK3* expression might also be altered in the brain of patients as MAPK phosphorylation levels in the amygdala were directly associated with anxiety symptoms in a previous study [77]. The authors demonstrate that the rate of extracellular signal-related kinase phosphorylation in the amygdala is negatively and independently associated with anxiety symptoms [77]. These findings further support our results of an involvement of MAPK signaling in SAD. Nevertheless, as BDNF is associated with several mental disorders, *MAPK3* differential expression may also not be restricted to social anxiety. An upregulation of MAPK-related genes is also found in Major Depressive Disorder (MDD, [78]) especially in cases with a history of ELA. Altered MAPK signaling involved in the development of mental disorders may be a result of stress on the mental or even cellular (e. g. infection) level. Indeed, early childhood adversities and infections are shown to severely affect the immune system resulting in an inflammatory phenotype and increase the risk for adult psychiatric disorders [9, 79]. Therefore, ELA might lead to molecular alterations mainly involved in inflammatory processes and those changes may induce an aberrant synaptic development in SAD transferred by MAPK signaling.

Gene co-expression analysis revealed gene clusters significantly associated with the emotional aspects of ELA (emotional abuse and neglect). Although no direct link on the level of differential gene expression was identified, this is an interesting finding as social anxiety and especially the emotional forms of childhood maltreatment are shown to significantly correlate [80, 81], which is supported by our data as well (Fig. S3). Therefore, a connection of emotional ELA and adult SAD on the molecular level seems likely. The MAPK3 interaction network shared one gene, *PTPN7*, with the red co-expression module, that significantly correlated with the scores of the CTQ subscales emotional abuse and emotional neglect as well as the total CTQ score. PTPN7 is a member of the phosphatase family and specifically inactivates MAPKs. Nothing is known about the regulation of PTPN7 in the context of ELA or SAD so far, although Schwieck et al. (2020) did not identify differential expression in MDD cases together with suicide risk and ELA history [78]. However, it acts as a regulator of MAPK signaling activity [82, 83]. The immune system is a plausible pathway how ELA could be molecularly involved in adult psychiatric disorders as ELA is known to cause inflammatory mimicking effects that are still measurable in adults, e.g. higher levels of typical markers of inflammation such as white blood cell count, circulating proinflammatory cytokine levels, and the acute phase molecule C-Reactive Protein (CRP) and lower NK cell activity [9, 84]. Although we did not measure cytokine levels or lower NK cell activity, none of the mentioned marker genes associated with emotional neglect and/or abuse were differing in the context of ELA and/or SAD in our sample (data not shown). Nevertheless, PTPN7 may be a promising candidate connecting ELA, the immune response and MAPK signaling as a potential regulator of anxiety disorders. Furthermore, blood cell type ratios were estimated and gene counts were adjusted to the ratios reflecting the immune activity. However, cell type ratios did not differ with respect to ELA or SAD in our sample.

The greenyellow co-expression module significantly correlated with emotional abuse and neglect in early childhood and was enriched for immune-related terms and JAK-STAT signaling. *STAT3* and *RAF1* were shared between this module and the MAPK3 ppi network. RAF1 is a known upstream regulator of MAPK signaling (Raf/MEK/ERK cascade, [85]). *RAF1* expression increases upon infection, which is mediated by interleukin 2 (IL-2, [86]), whereas inhibition of *RAF1* affects production of IL-6 and IL-8 in cultured human corneal epithelial cells [87]. STAT3—a transcription factor—regulates processes involved in inflammation and tumorigenesis by regulating cell proliferation, differentiation and metabolism [88]. STAT3 is a member of the JAK-STAT signaling pathway whose canonical mode is based on cytokine release followed by MAPK signaling activation [89]. In the non-canonical signaling pathway, unphosphorylated STATs are localized on heterochromatin in the nucleus in association with proteins regulating the maintenance of heterochromatin state [90]. Therefore, STAT3—like MAPK3 and RAF1, respectively—is involved in a complex crosstalk of signaling pathways and may be involved in epigenetic regulation of downstream processes. A *STAT3* knockout in mice leads to reduced negative behavioral reactivity [91] Additionally, *STAT3* is involved in alcohol withdrawal [92] and depressive symptoms in rats [93, 94]. Therefore, RAF1 and STAT3 are potential candidates connecting ELA, immune response and SAD.

Gene set functional enrichment of the genes overlapping in the turquoise module, that significantly correlated with ELA, and the MAPK3 ppi network revealed mainly terms related to signal transduction pathways (i.e. MAPK and NTRK and neurotrophin signaling, Fig S10). Amongst others, NTRK signaling was enriched, pointing towards a role of those genes in BDNF-related processes. Moreover, the gene set enrichment of the MAPK3 interactome substantiates the role of NTRK signaling in the association of ELA and SAD.

## Summary and model

As stated above, ELA is known to have an effect on neuronal structures by affecting the immune system (e. g. cytokine levels), which is linked to structural changes involved in the development of mental disorders. Our findings support this assumption on the molecular level: The gene set enrichment of the co-expression clusters revealed terms especially involved in the cellular structure, signal transduction and immune response. Genes co-expressed in clusters associated especially with emotional ELA are potential interactors of *MAPK3*, which is significantly differentially expressed in individuals suffering from SAD compared to controls. Especially *STAT3*, the hub gene of such a co-expression cluster, may be regulated by the emotional ELA-dependent release of interleukins like IL-6 [95, 96] and thus may be involved in the cell type-specific regulation of more growth factor and cytokine release [97-99] which for their part increase *MAPK3* expression [100]. *MAPK3* may be further involved in the expression of genes shaping synaptic plasticity, e.g. *BDNF*.

In a recent study of our group, the blood DNA methylome was analyzed in the same cohort presented here. Differentially methylated regions (DMRs) specifically associated with SAD, ELA, or the interaction of SAD and ELA were identified [101]. None of these regions were overlapping with the DEGs found in the current gene expression analyses. However, STAT3 is shown to interact with the DNA methyltransferase DNMT1 [102, 103]. Therefore, STAT3-directed DNA methylation is a possible step in the signal transduction cascade transferring ELA to SAD. In the study of Camilo *et al*. (2019), a gene network approach revealed a direct association of *MAPK3* methylation and cocaine use disorder [104]. Therefore, in a follow-up study, we aim to conduct a multivariate machine learning-based analysis to integrate DNA methylation and gene expression data in the context of SAD following ELA.

The approach presented here has several limitations: First, the expression of genes can vary between blood cell types and therefore, differential cell type composition can affect results. Immune responsive cell types are known to play a role in mental disorders such as anxiety-related MDD or panic disorder [105-107], which make them unsuitable for usage as covariate in statistical tests and DGE analysis. Therefore, we adjusted the gene expression data to the estimated cell type composition with the help of reference cell counts of a subset of our cohort and showed that the number of real counts and estimated ratios correlated for several relevant cell types (Fig. S1). Furthermore, we have to be aware of the fact that transcriptomic profiles are not only cell type-but also tissue-specific, and that we therefore cannot assume that the differences we observe in blood directly reflect the situation in brain (as mentioned for *BDNF* earlier). In psychiatric transcriptomics, we are faced with the problem, that the tissue of interest— the brain—is not easily available for transcriptomic analyses in living individuals. However, we can assume that there is some overlap of genes expressed similar in blood and brain, as human whole blood tissue showed a significant similarity in gene expression to multiple brain tissues with a median correlation of 0.5 as revealed by microarray analysis [108]. Furthermore, rat brain and blood tissue more than half of the 29,215 genes analyzed by microarray were co-expressed [109]. Furthermore, age differed significantly between the groups with high and low levels of ELA. Age effects were not identified for the expression of the candidate genes (Fig. S3) and co-expression clusters correlating with age were excluded from further analyses. An approach in a larger cohort would be needed to decipher whether age-dependent gene expression has an effect on the results presented here. Moreover, the phosphorylation levels of MAPK3 and MAPK signaling (and therefore the activation of the signal transduction cascade) mark an important step in the signal cascade that should be included in future experiments to clarify whether gene product or phosphorylation abundance are the potential drivers behind the molecular development of SAD.

In summary, by investigating gene expression in context of SAD and its relation to ELA on a transcriptome level, we were able to identify DEGs associated with SAD—with *MAPK3* being the most significant DEG—as well as co-expression clusters correlating with ELA and/or its subclasses. Interestingly, functional enrichment of MAPK3 protein-protein interaction network and ELA associated gene co-expression modules pointed towards signal transduction pathways and the immune system. Additionally, shared genes are involved in JAK-STAT and ERK signaling as well as DNA methylation. Although a direct molecular link of ELA leading to adult SAD by gene expression changes was not identified, the data indicate an indirect relation of emotional ELA and SAD mediated by the interaction of genes involved in immune-related signal transduction. Further studies will be needed to replicate our findings in independent, larger cohorts and to investigate the potential effect of the immune responsive gene expression pattern caused by ELA on adult anxiety disorders in more detail.

## 6. Data availability statement

The raw data supporting the conclusions of this article will be made available by the authors, without undue reservation.

## 7. Ethics statement

The studies involving human participants were reviewed and approved by Ethics committee of the University of Tuebingen (process number 526/2018BO2). The participants provided their written informed consent to participate in this study.

## 9. Acknowledgments

We acknowledge the support by the NGS Competence Center Tuebingen. The authors also wish to express their appreciation to all participants.

