## Supplemental Figures for "Blood transcriptome analysis suggests an indirect molecular association of early life adversities and adult social anxiety disorder by immune-related signal transduction"

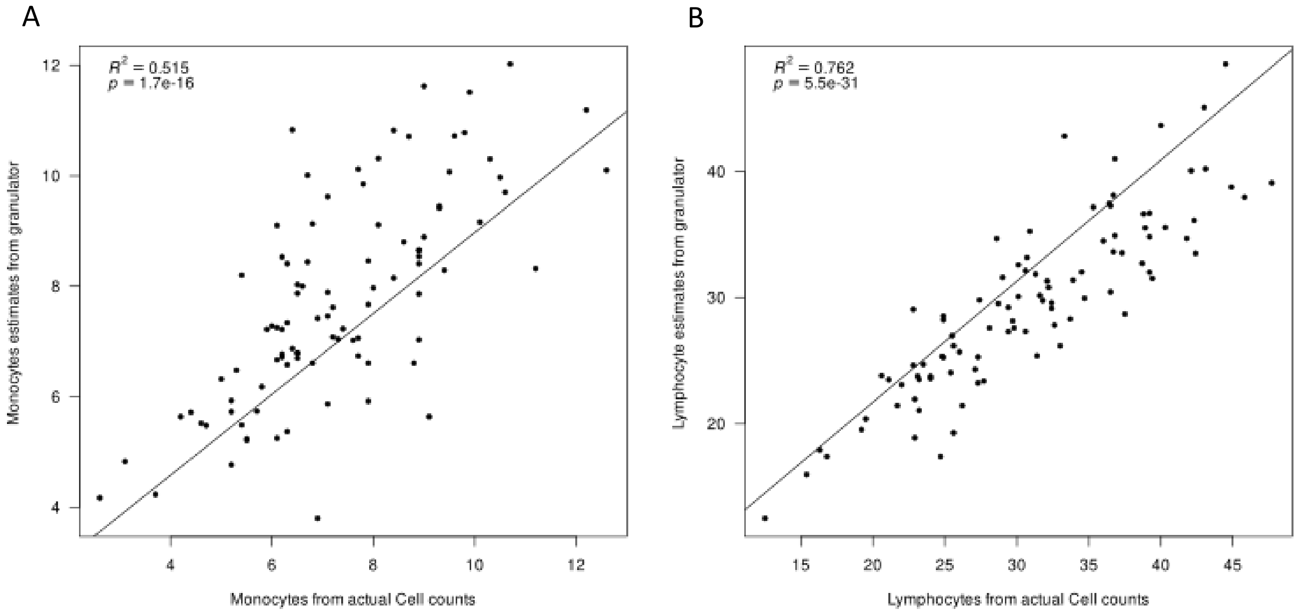


**Figure S1**: Correlations of cell type ratios estimated using *granulator* and real counts from a cohort subset for **A** Monocytes and **B** Lymphocytes.

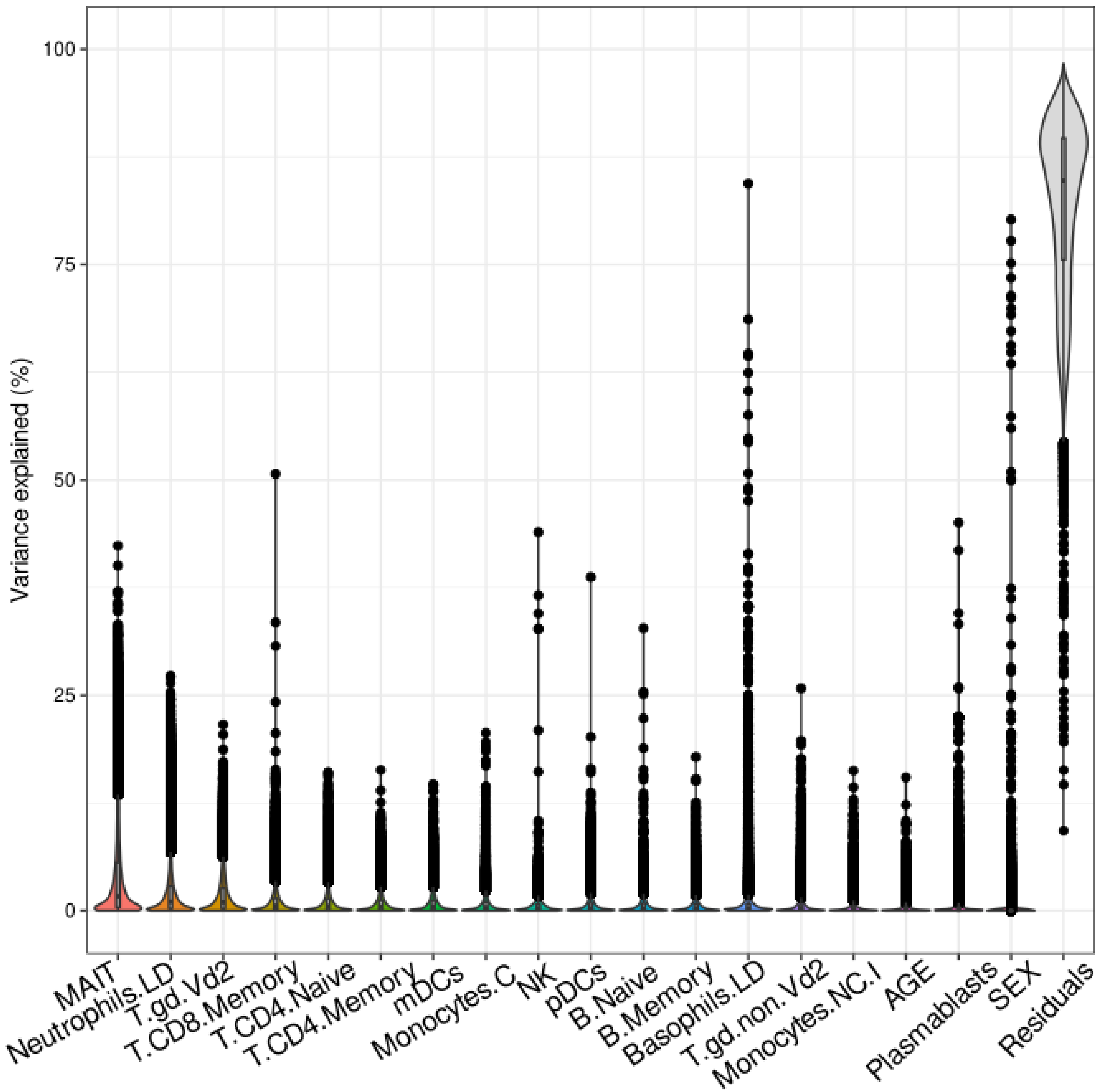


**Figure S2**: Explained variance by variables in normalized gene expression prior to gene count adjustment.

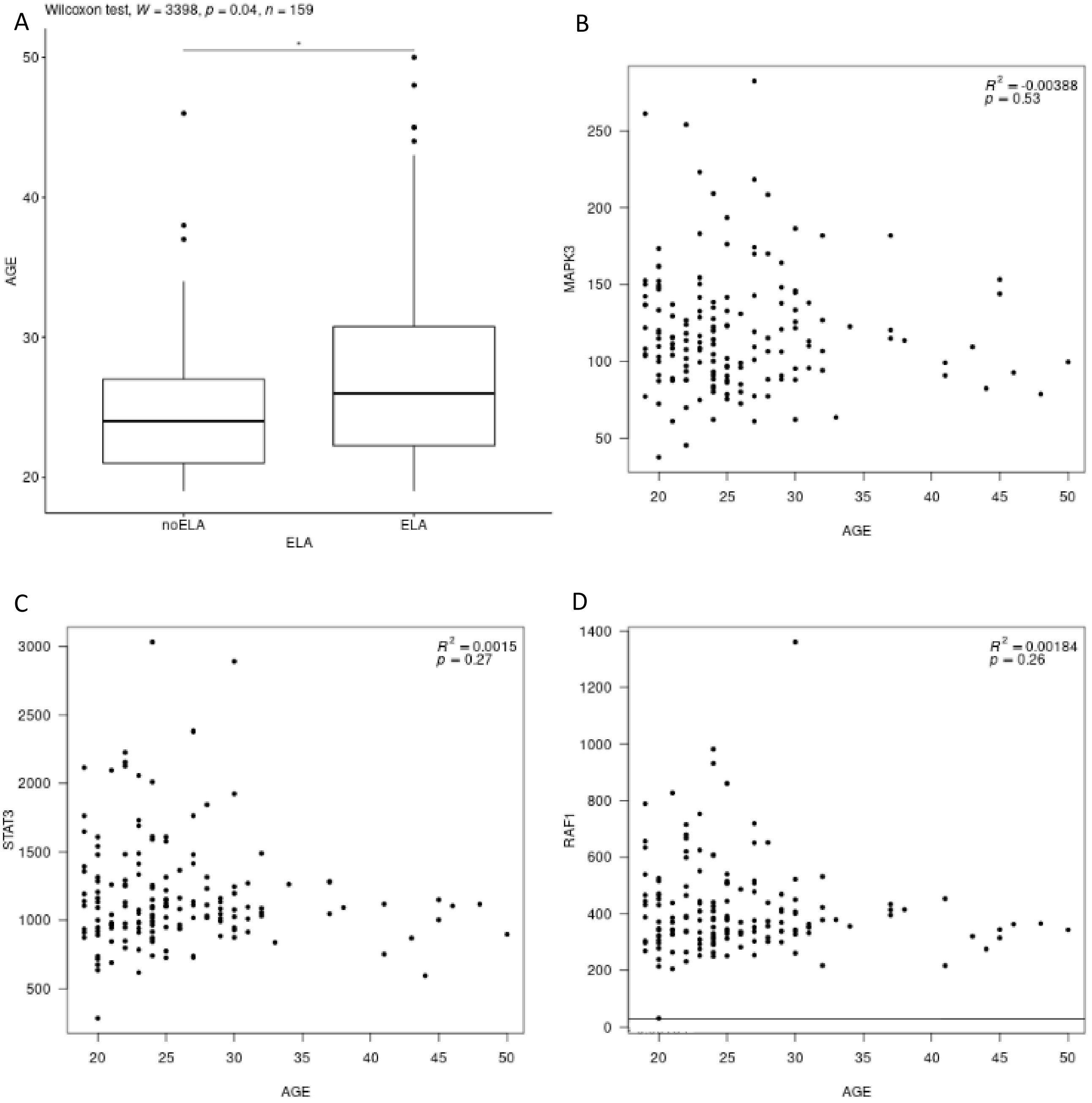


**Figure S3**: **A** The age distribution is shown among the groups with high and low levels of ELA, respectively. However, there is no correlation of age and gene expression of representative genes, namely **B** *MAPK3*, **C** *STAT3* and **D** *RAF1*.

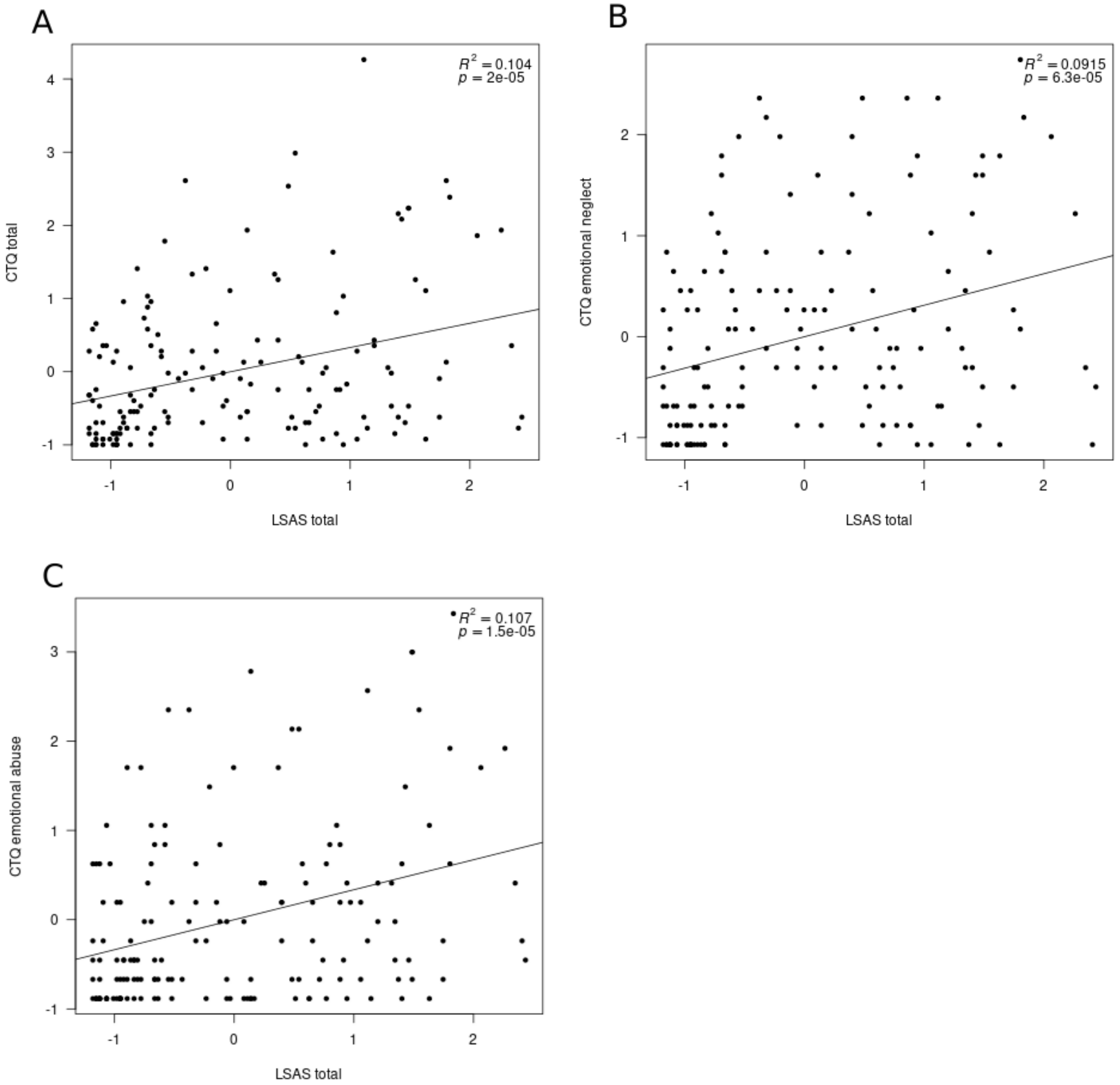


**Figure S4**: Correlations of scaled CTQ and LSAS score with **A** CTQ total, **B** emotional neglect and **C** emotional abuse.

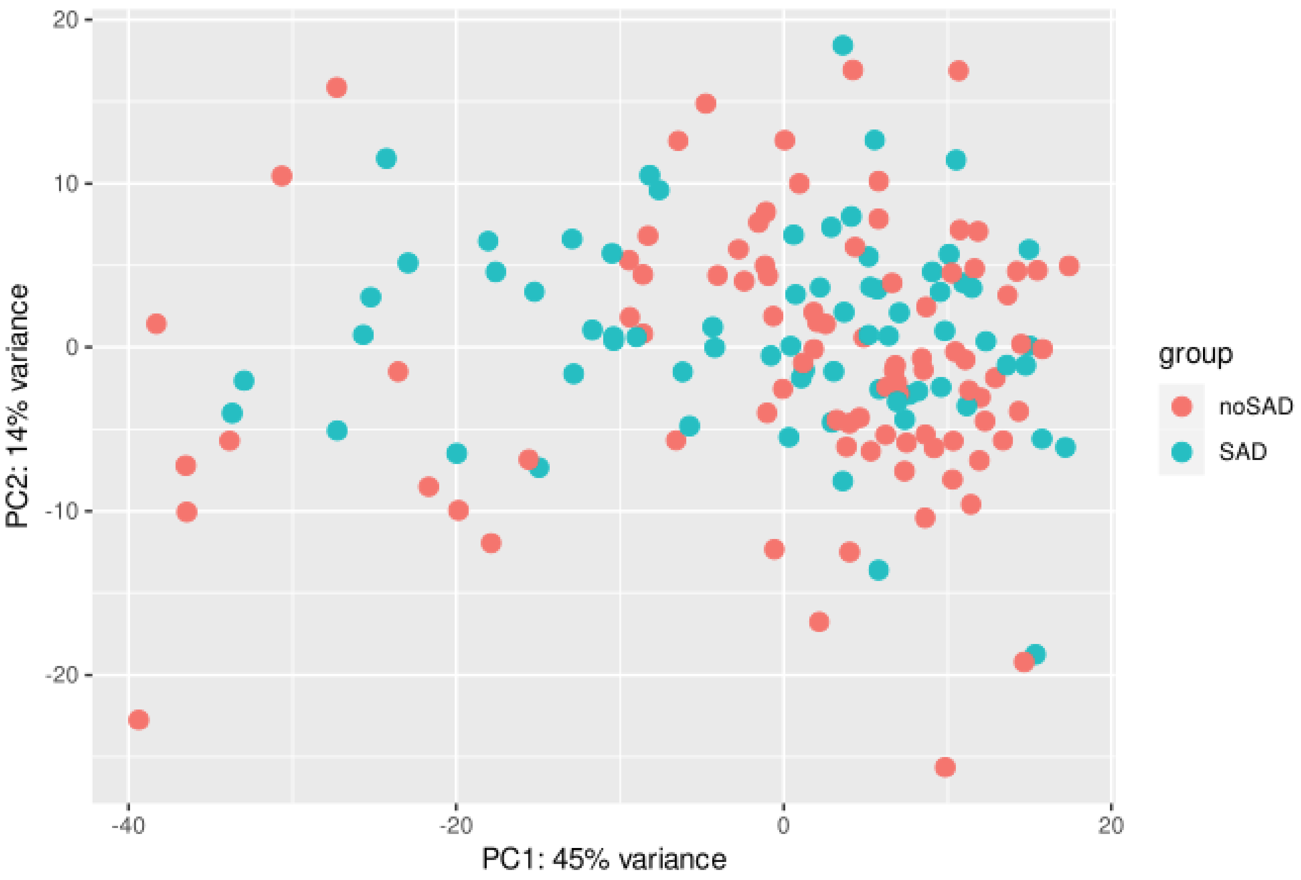


**Figure S5**: Principal component analysis of normalized and cell type ratio adjusted gene expression data with respect to SAD.

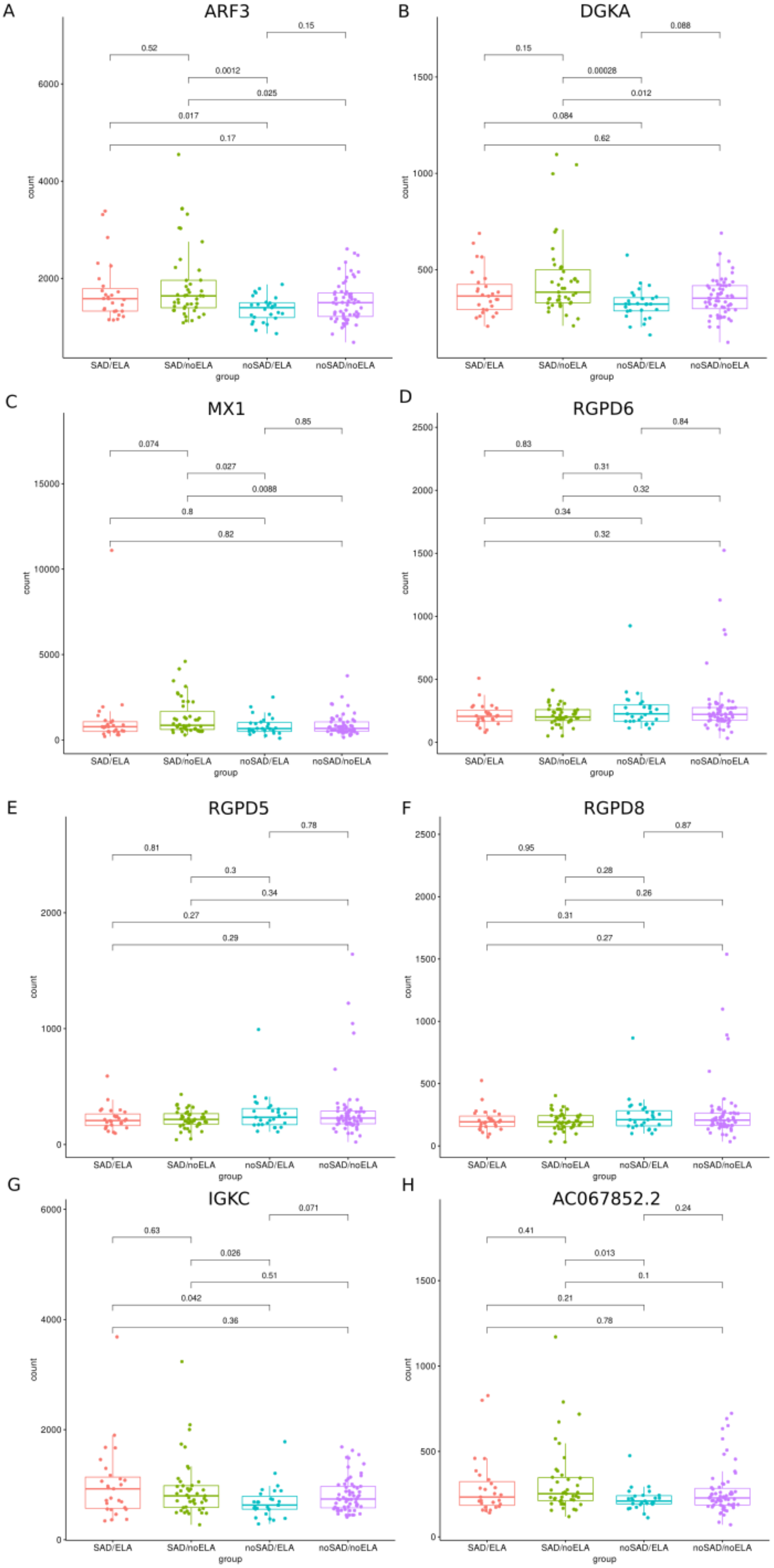
A
ARF3
0.15
0.52
0.0012
0.025
0.017
0.17
count
group
SAD/ELA
SAD/noELA
noSAD/ELA
noSAD/noELA
B
DGKA
0.088
0.15
0.00028
0.012
0.084
0.62
C
MX1
0.85
0.074
0.027
0.0088
0.8
0.82
D
RGPD6
0.84
0.83
0.31
0.32
0.34
0.32
E
RGPD5
0.78
0.81
0.3
0.34
0.27
0.29
F
RGPD8
0.87
0.95
0.28
0.26
0.31
0.27
G
IGKC
0.071
0.63
0.026
0.51
0.042
0.36
H
AC067852.2
0.24
0.41
0.013
0.1
0.21
0.78

**Figure S6**: Gene expression patterns of the eight significantly differentially expressed genes (DEGs) between the SAD group and controls that exhibited an expression rather caused by outliers or other effects than being a true DEG. The counts are shown with respect to SAD and ELA. The Wilcoxon rank sum test was applied and the p values were adjusted for multiple testing using Benjamini-Hochberg correction. Expression patterns are shown for **A** ADP-ribosylation factor 3 (*ARF3*), **B** Diacylglycerol Kinase Alpha (*DGKA*), **C** MX Dynamin Like GTPase 1 (*MX1*), **D** RANBP2 Like And GRIP Domain Containing 6 (*RGPD6*), **E** RANBP2 Like And GRIP Domain Containing 5 (*RGPD5*), **F** RANBP2 Like And GRIP Domain Containing 8 (*RGPD8*), **G** Immunoglobulin Kappa Constant (*IGKC*) and **H** the long non-coding RNA *AC067852.2*.

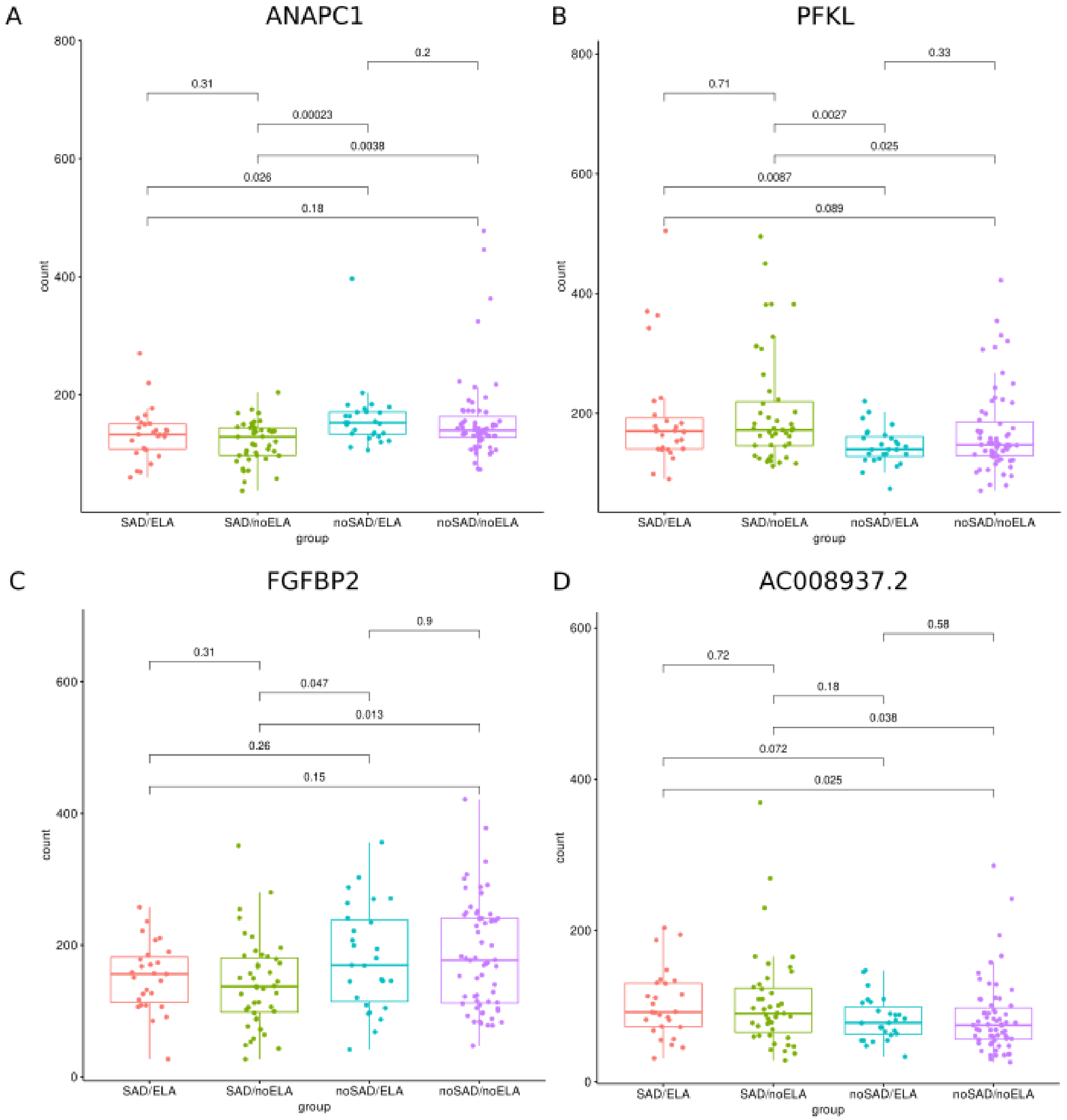


**Figure S7**: Gene expression patterns of four significantly differentially expressed genes (DEGs) between the SAD group and controls showing a clear SAD associated expression pattern. The counts are shown with respect to SAD and ELA. The Wilcoxon rank sum test was applied and the p values were adjusted for multiple testing using Benjamini-Hochberg correction. Expression patterns are shown for **A** Anaphase Promoting Complex Subunit 1 (*ANAPC1*), **B** Phosphofructokinase, Liver type (*PFKL*), **C** Fibroblast Growth Factor Binding Protein 2 (*FGFBP2*) and **D** the long non-coding RNA *AC008937.2*.

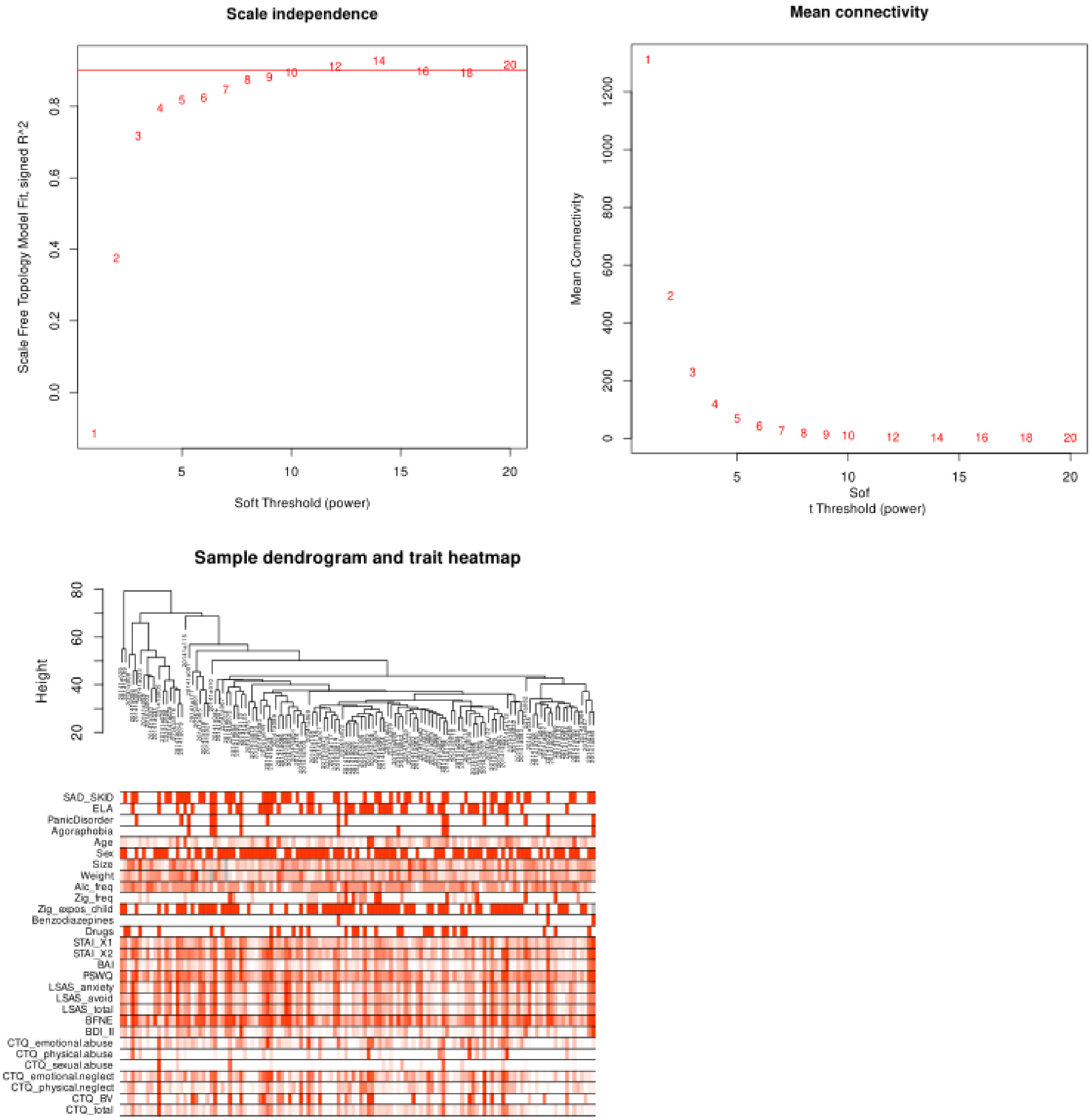


**Figure S8**: Weighted Gene Co-expression Analysis (WGCNA). **A** Selection of the soft threshold by Scale-free topology fitting index R2 analysis and **B** mean connectivity for various soft threshold powers. **C** Sample dendrogram and trait heatmap revealed no severely deviating samples.

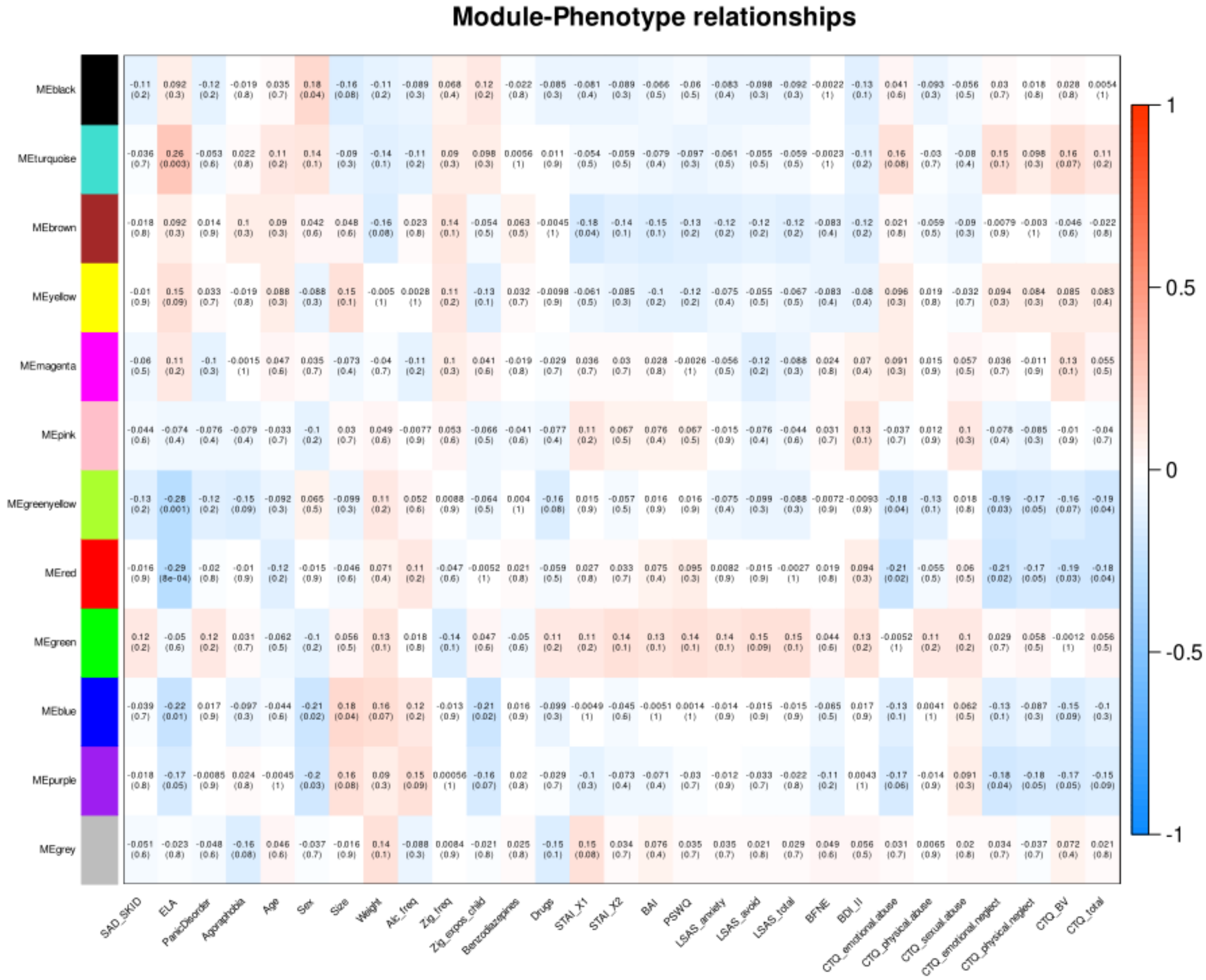


**Figure S9**: Heatmap of the correlation between the variables and module eigengenes. Correlation coefficients are followed by the *p* value in brackets.

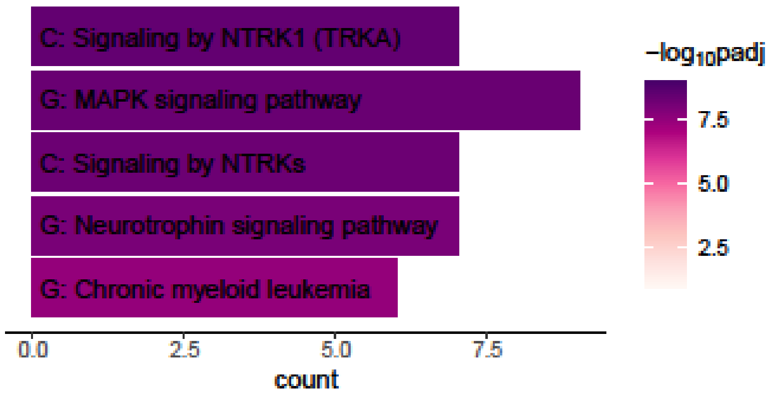


**Figure S10:** Gene functional enrichment of genes overlapping between the MAPK3 ppi String network and the turquoise *WGCNA* co-expression module.
